# Complete genome sequences of two new strains of the shipworm endosymbiont, *Teredinibacter turnerae*

**DOI:** 10.1101/2024.08.13.607755

**Authors:** Mark T. Gasser, Ron Flatau, Marvin Altamia, Claire Marie Filone, Dan Distel

**Affiliations:** Johns Hopkins University Applied Physics Laboratory, Laurel, Maryland, USA 20723; Ocean Genome Legacy Center, Northeastern University, Nahant, Massachusetts, USA 01908

## Abstract

We present the complete genome sequences of two strains of *Teredinibacter turnerae*, SR01903 and SR02026, shipworm endosymbionts isolated from the gills of *Lyrodus pedicellatus* and *Teredo bartschi*, respectively, and derived from Oxford Nanopore sequencing. These sequences will aid in the comparative genomics of shipworm endosymbionts and understanding of host-symbiont selection.

## Announcement

*Teredinibacter* species are cultivable intracellular symbionts of xylotrophic, bivalve wood-borers (Teredinidae) (1–4) and have been shown to secrete lignocellulolytic enzymes that aid in host digestion (5, 6). Wood containing live specimens of *Lyrodus pedicellatus* and *Teredo bartschi* was collected from the Indian River Lagoon, Merit Island, FL. (N 28.40605 W 80.66034) on January 24, 2020, and subsequently maintained in laboratory culture. Strain SR01903 was isolated from the gill of a single specimen of *L. pedicellatus* immediately after collection from the wild. Strain SR02026 was isolated from the gill of a fourth-generation lab-reared specimen of *T. bartschi*. Bacterial isolations were performed as in O’Connor *et al*., 2014. Briefly, gills were removed by dissection and homogenized in 1.0 mL of SBM medium (7) in an autoclave-sterilized glass dounce homogenizer. Homogenates were streaked onto culture plates containing 1.0% Bacto agar prepared in shipworm basal medium (SBM) at pH 8.0 supplemented with 0.2% w/v powdered cellulose (Sigmacell Type 101; Sigma-Aldrich) and 0.025% NH_4_Cl.

Plates were then incubated at 30 °C. When individual colonies appeared, a single colony was picked, re-streaked, and regrown. This process was repeated until clonal isolates were achieved. Genomic DNA was extracted from the resulting clonal isolates, as in O’Connor *et al*., 2014 using the Qiagen DNeasy Blood and Tissue Kit following the manufacturer’s recommended protocol for cultured cells with the exception that DNA was eluted with two 75 μL volumes of AE buffer preheated to 56 °C. DNA quality and length were assessed on Tapestation (Agilent Technologies, US). Nanopore (Oxford Nanopore Technologies, UK) sequencing was performed without DNA fragmentation or size selection. The sequencing library was prepared using the Q20+ chemistry Ligation Sequencing Kit (SQK-LSK112) and sequenced on a MinION (Mk1B) instrument using a R10.4 (FLO-MIN112) flow cell. Bases were called using Guppy v6.4.6 with the high-accuracy (HAC) algorithm, and default read quality filtering. Adapters were trimmed from reads using Porechop v0.2.4 (https://github.com/rrwick/Porechop) and filtered to remove reads less than 1 Kb using Filtlong v0.2.1 (https://github.com/rrwick/Filtlong). De novo assembly was performed with Flye v2.9.2 (https://github.com/fenderglass/Flye) (8) followed by contig correction and consensus generation with Racon v1.5.0 (https://github.com/lbcb-sci/racon) and Medaka v1.8.0 (https://github.com/nanoporetech/medaka). Assemblies were then circularized and rotated to start at *dnaA*, predicted by prodigal v2.6.3 (9) with Circlator v1.5.5 (https://github.com/sanger-pathogens/circlator) (10). Chromosomal assemblies were produced for both strains and annotated using the NCBI Prokaryotic Genome Annotation Pipeline (PGAP) (Table 1). All software was run using default settings unless otherwise noted. The primary sequences were 98.67% identical based on the calculated average nucleotide identity (ANI) (11) and highly syntenic (Fig. 1) (12).

**Table 1.** Genome sequencing, assembly, and annotation of Teredinibacter strains.

| Strain | SR01903 | SR02026 |
| --- | --- | --- |
| Host | *L. pedacellatus* | *T. bartschi* |
| Reads | 77,041 | 253,198 |
| Bases (M) | 289.9 | 306.2 |
| Read N50 | 11,088 | 2,331 |
| Assembly (bp) | 5,229,132 | 5,163,139 |
| Coverage | 52x | 44x |
| GC content (%) | 50.9 | 50.9 |
| Genes | 4,207 | 4,142 |
| CDS | 4,152 | 4,086 |
| rRNA (complete) | 3 | 3 |

**Figure 1.**
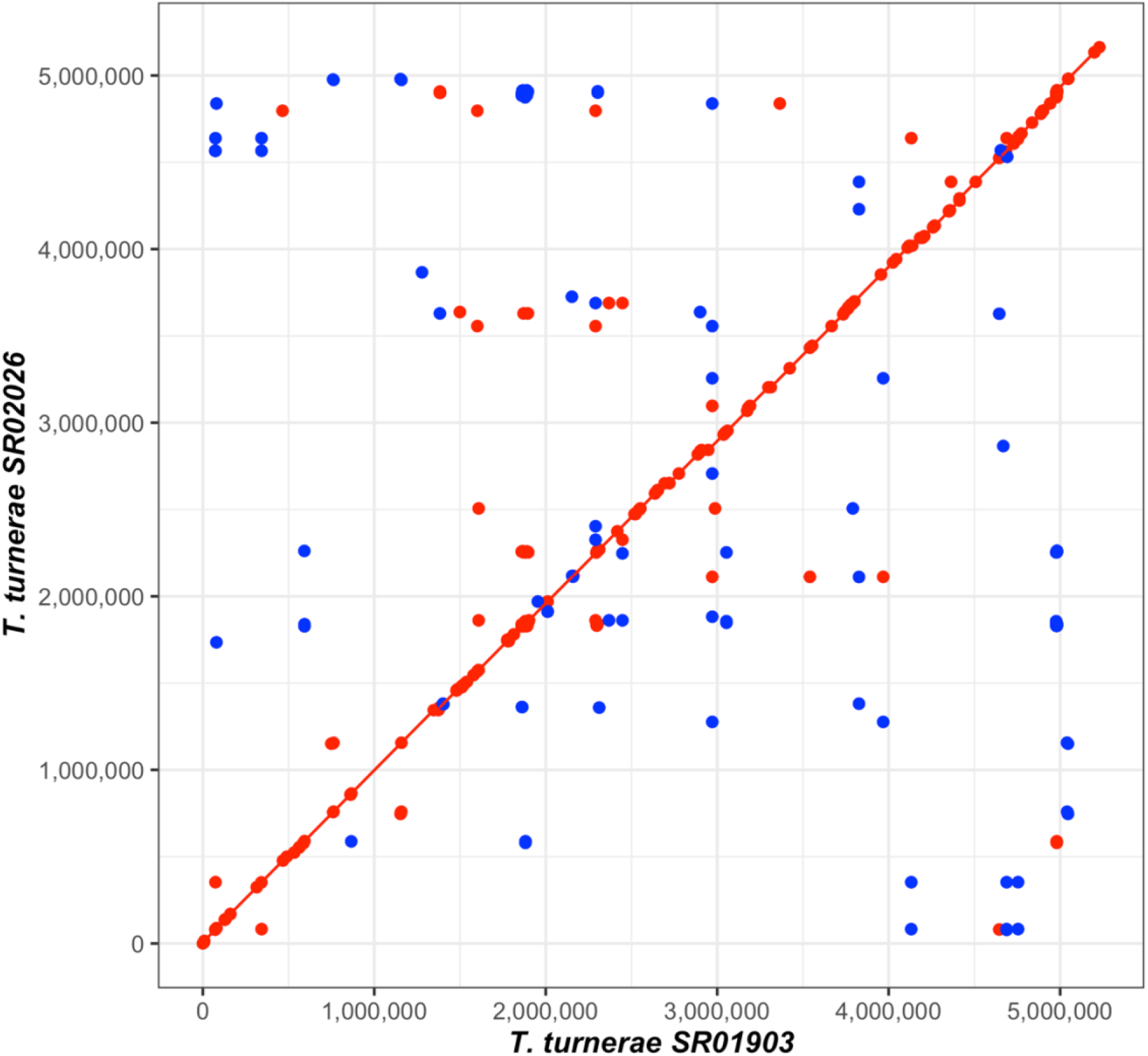
Synteny plot comparing the genome sequences of SR01903 and SR02026. A MUMmer3 plot was generated with NUCmer v3.1 (12) using NUCmer to assess synteny and completion. Minimum exact matches of 20bp are represented as a dot with lines representing match lengths >20bp. Forward matches are displayed in red, while reverse matches are shown in blue.

## Data availability

The complete genome sequences for SR01903 and SR02026 have been deposited in GenBank under the accession numbers CP149818 and CP149819, respectively. The Oxford Nanopore sequencing reads are available from the NCBI Sequence Read Archive (SRA) under the accession numbers SRR28421271 and SRR28421270, respectively.

## Acknowledgments

Research reported in this publication was supported by the following awards to DLD: National Oceanic and Atmospheric Administration (NA19OAR0110303), Gordon and Betty Moore Foundation (GBMF 9339), National Institutes of Health (1R01AI162943-01A1, subaward: 10062083-NE), and Johns Hopkins University Applied Physics Laboratory internal research and development funds. The National Science Foundation (DBI 1722553) also funded some equipment used in this research. The funders had no role in study design, data collection and interpretation, or the decision to submit the work for publication.

